# Information-based summary statistics for spatial genetic structure inference

**DOI:** 10.1101/2021.08.31.458463

**Authors:** Xinghu Qin, Oscar E. Gaggiotti

## Abstract

1. Inference of spatial patterns of genetic structure often relies on parameter estimation and model evaluation using a set of summary statistics (SS) that summarise the information present in the data. An important subset of these SS is best described as diversity indices, which are based on information theory principles that can be classified as belonging to three different ‘families’ encompassing a spectrum of information measures, ^*q*^*H*. These include the richness family of order *q* = 0, ^*Ar*^*SS*; the Shannon family of order *q* = 1, ^*H*^*SS*; and the heterozygosity family of order *q* = 2, ^*He*^*SS*. Although commonly used by ecologists, the Shannon family has been rather neglected by population geneticists and evolutionary biologists. However, recent population genetic studies have advocated their use, yet the power of these SS for spatial structure discrimination has not been systematically assessed.
2. In this study, we performed a comprehensive assessment of the three families of SS, as well as a fourth family consisting of SS belonging to the Shannon family but expressed in terms of Hill numbers 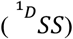, for spatial structure inference using simulated microsatellites data under typical spatial scenarios. To give an unbiased evaluation, we used three machine learning methods, Kernel Local Fisher discriminant analysis (KLFDA), random forest classification (RFC), and deep neural network (DL), to test the performance of different SS to discriminate between spatial scenarios, and then identified the most informative metrics for discriminatory power.
3. Results showed that the SS family of order *q* = 1 expressed in terms of Hill numbers, 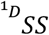, outperformed the other two families (^*Ar*^ *SS*, ^*He*^ *SS*) as well as the untransformed Shannon entropy (^*H*^ *SS*) family. Jaccard dissimilarity (*J*) and its Mantel’s *r* showed the highest discriminatory power to discriminate all spatial scenarios, followed by Shannon differentiation Δ*D* and its Mantel’s *r*.
4. Information-based summary statistics, especially the diversity of order *q* = 1 and Shannon differentiation measures, can increase the power of spatial structure inference. In addition, different sets of SS provide complementary power for discriminating between spatial scenarios.

## Introduction

Spatial biodiversity patterns generated by different evolutionary and demographic processes can be observed at the ecological or species level and, at the genetic or molecular level (Van Tienderen 1991; Novembre & Stephens 2008; Fortuna *et al*. 2009; Wang *et al*. 2011; Stotz, Gianoli & Cahill 2016). However, metrics and approaches to describe these spatial patterns and to infer the underlying processes differ greatly between these two biodiversity levels. The metrics used to study ecological variation (species) and genetic variation (alleles) are mainly dominated by the traditional indices in their own domains, such as species richness, Shannon index in ecology, and allelic richness, heterozygosity in population genetics. These indices comprise a spectrum of information measures (*q* profile, ^*q*^*H*; (Hill 1973; Jost 2006)), including richness (*q*=0, *S*), Shannon entropy (*q*=1, *H*), and heterozygosity (or Gini-Simpson index, *q*=2, *He*). Each index of order *q* provides a different type of information, with the index of order *q=*0 emphasising rare elements, the index of order *q* = 2 emphasising common elements, while the index of order *q*=1 measuring uncertainty in proportion to their frequency, neither preferring rare elements nor common elements (Sherwin *et al*. 2017). However, *q* =1 family only received sporadic attention in population genetics.

The use of summary statistics has facilitated our understanding of ecology and evolution in terms of describing spatial biodiversity patterns (e.g., Distance-Decay (Nekola & White 1999)), and examining likely processes underlying them. Typically, this is done by decomposing total diversity (*γ*-diversity) into within-aggregate (*α*-diversity) and between aggregates (*β-*diversity) based on species’ or community spatial aggregation (Lande 1996; Ricotta 2005). The derived *β-*diversity is then used to examine the dissimilarity or differentiation between aggregates. Two main decomposition methods have been used to do this, multiplicative (*SS*_*γ*_ =*SS*_*β*_ *\*SS*_*α*_) and additive (*SS*_*γ*_=*SS*_*β*_ + *SS*_*α*_) decomposition (Ricotta 2005). The desirable *β-*diversity should be additive when pooling or partitioning the aggregates and should represent the actual proportion of non-shared elements (true dissimilarity or differentiation) due to divergence or differentiation between aggregates (Chao, Chiu & Jost 2014). *β-*measures additively decomposed from *q*=0 (e.g., Jaccard dissimilarity) don’t measure true dissimilarity, because they only count presence and absence ignoring the abundance or frequency of elements. *β-*measures of order *q* = 2, the well-known fixation index, *F*_*ST*_ family (including *F*_*ST*,_ *G*_*ST*_ etc.) derived from the multiplicative decomposition of heterozygosity (*He*), don’t measure true differentiation (true dissimilarity) and are not independent of *α*-diversity or *γ*-diversity (Jost 2008; Ma, Ji & Zhang 2015). On the other hand, *β-*measures based on Shannon entropy (order *q* = 1, the Shannon differentiation, Δ*D*), measure true differentiation and satisfy monotonicity without dependence problem (Gaggiotti *et al*. 2018), which are desirable metrics for measuring differentiation.

A common difficulty faced when measuring biodiversity with standard metrics is that, with the exception of richness, they do not have an intuitive interpretation in terms of the number of effective elements in the system (Jost 2006). However, this problem is easily overcome by using Hill numbers (Hill 1973), and this is the approach we use in the present study. Thus, allelic richness is represented by ^0^*D* while the effective number of alleles based on Shannon entropy and heterozygosity are given by ^1^*D* and ^2^*D* respectively.

Diversity at one level of biological organization (community, species) may sustain the diversity at the other (Lankau & Strauss 2007). Thus, in addition to describing diversity patterns, researchers have made substantial efforts to unify the two levels of biodiversity (species diversity of ecological communities and genetic diversity of populations) and to reveal ecological and evolutionary processes underpinning their spatial patterns (Vellend 2005). However, these so-called Species-Genetic-Diversity-Correlation (SGDC) studies have rarely measured the two types of diversity consistently (Gaggiotti *et al*. 2018). Integrative studies of species and genetic diversity, and the ecological factors underlying their association or lack thereof using the same type of index would contribute to a better understanding of eco-evolutionary dynamics.

The use of informative diversity metrics is crucial, not only for detecting changes in biodiversity patterns but also for understanding the demographic and evolutionary history of species (Csilléry *et al*. 2010). The performance of population genetics summary statistics has been thoroughly evaluated in the context of spatial demographic inference (Alvarado-Serrano & Hickerson 2016) and similar studies are needed for equivalent statistics based on Shannon entropy. The present study represents the first step in this direction by evaluating the power of the information-based diversity measures (represented by ^1^*D* and Shannon differentiation, Δ*D*) and comparing it with that of traditional measures (represented by allelic richness, heterozygosity, and their *β-*diversity measures) to discriminate between spatial scenarios using recent state-of-the-art machine learning approaches.

We simulated microsatellite data under five spatial scenarios that include panmixia, finite island model, hierarchical island model, stepping-stone model and hierarchical stepping-stone model, which are the typical spatial demographic models that have been used to describe the spatial structure of natural populations in fragmented landscapes. We employed three state-of-the-art machine learning approaches, kernel local discriminant analysis (KLFDA), conditional random forest classification, and deep neural networks (*DNN*) to characterize the behaviour of these diversity metrics for discriminating different spatial scenarios. Our results showed that information-based summary statistics can provide more power than traditional measures to make inferences about spatial genetic structure.

## Methods

To evaluate the ability of the new SS and traditional SS in discriminating different spatial scenarios, we simulated five spatial scenarios that encompass hierarchical and non-hierarchical population structures using coalescent simulations. More specifically, we considered populations without hierarchical structure and populations structured into three hierarchical levels, ecosystem, aggregate (e.g., region) and sub-aggregate (e.g., population) level.

We calculated the traditional and new summary statistics from these scenarios and then used the state-of-the-art machine learning approaches to test their power to discriminate among spatial scenarios.

### Models and model parameters

We considered five spatial scenarios, panmixia, island model, hierarchical island model, stepping-stone model, and hierarchical stepping-stone model. Instead of using fixed values for the parameters, we sampled them from probability distributions. Table 1 presents all scenarios and the respective parameter distributions used in the simulations. For the island model, stepping-stone model, hierarchical island model and hierarchical stepping-stone model, each scenario consisted of 16 populations with population size sampled from *U*(100,1000). For the panmixia model, we simulated one panmictic population, with population size drawn from *U*(1600,16000). The hierarchical island models consist of four regions with each region comprising 4 populations. In terms of the hierarchical stepping-stone models, we simulated two regions with each region comprising 8 populations. We assume a stepwise mutation model with a constant mutation rate of 5×10^−4^ for all scenarios. In the case of the non-hierarchical scenarios (island model and stepping-stone model), the migration rate, *m*, was drawn from a uniform distribution *U*(0.001,0.1). In the case of the hierarchical scenarios, migration rates between pairs of populations within regions were sampled from *U*(0.001,0.1) and migration rates between populations from different regions were sampled from *U*(0.00005,0.005).

**Table 1.** Parameters used in the simulations. In the case of the panmixia scenario, we simulated a single population but generated 16 samples at random. In the case of the hierarchical models, we indicate the number of populations per region in parenthesis.

| Scenarios | Regions | Number of populations | Population size | Sample size | Migration rate | Mutation rate | Number of loci |
| --- | --- | --- | --- | --- | --- | --- | --- |
| Panmixia | 1 | 1 (16) * | $U(1600, 16000)$ | 320 | | $5 \times 10^{-4}$ | 10 |
| Island model | 1 | 16 | $U(100, 1000)$ | 20 | $U(0.001, 0.1)$ | $5 \times 10^{-4}$ | 10 |
| Hierarchical island model | 4 (4,4,4,4) | 16 | $U(100, 1000)$ | 20 | $m_{\text{within}}: U(0.001, 0.1)$<br>$m_{\text{between}}: U(5E-5, 5E-3)$ | $5 \times 10^{-4}$ | 10 |
| Stepping-stone | 1 | 16 | $U(100, 1000)$ | 20 | $U(0.001, 0.1)$ | $5 \times 10^{-4}$ | 10 |
| Hierarchical stepping-stone | 2 (8,8) | 16 | $U(100, 1000)$ | 20 | $m_{\text{within}}: U(0.001, 0.1)$<br>$m_{\text{between}}: U(5E-5, 5E-3)$ | $5 \times 10^{-4}$ | 10 |

### Simulations

The coalescent-based simulator fastsimcoal2 (Excoffier & Foll 2011; Excoffier *et al*. 2013) was used to generate microsatellite synthetic data under the five scenarios described above. For each of these five spatial scenarios, we simulated 10 independent microsatellite loci sharing the same mutation rate. 100 sets of parameters (100 simulations) were randomly drawn from prior distributions, and each parameter set was used to generate 1000 replicate data sets. We sampled 20 individuals per population under each spatial model (standard and hierarchical versions of the island and stepping-stone models). In the case of the panmixia model, we sampled 320 individuals and then randomly partitioned them into 16 samples consisting of 20 individuals each to obtain a set of samples equivalent to those of the other four scenarios.

### Summary Statistics

We chose the commonly used genetic diversity indices, allelic richness (*Ar*, noted ^*Ar*^*SS* hereafter) and heterozygosity (*He*, noted ^*He*^*SS* hereafter) as well as their corresponding *β*-diversity measures, Jaccard dissimilarity (Jaccard 1912) and fixation index (Weir & Cockerham 1984), as the traditional summary statistics. The allelic richness and expected heterozygosity were partitioned into three hierarchical levels, population level, regional level and ecosystem level, with the corresponding measures being, 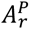 (allelic richness at the population level), 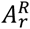 (allelic richness at the regional level), 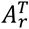 (total allelic richness in the ecosystem) and 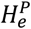 (expected heterozygosity at the population level), 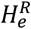 (expected heterozygosity at the regional level), 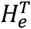 (total heterozygosity in the ecosystem). Accordingly, the *β*- measures were partitioned into 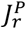 (Jaccard dissimilarity among populations within a region) and 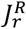 (Jaccard dissimilarity among regions within an ecosystem) for allelic richness, and 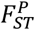 (*F*_*ST*_ among populations within a region) and 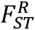 (*F*_*ST*_ among regions within an ecosystem) for expected heterozygosity.

We chose the diversity of order *q*=1, the transformed Shannon “effective number”-^1^*D*, as well as Shannon differentiation (Δ*D*) as the new summary statistics 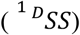. ^1^*D* was also decomposed into population level, regional level and ecosystem level, which were *D*_*γ*_, *D*_*α*_^2^, *D*_*α*_^1^, respectively. The equivalent number of regions and the equivalent number of populations thus were *D*_*β*_^2^, *D*_*β*_^*1*^, respectively. In the same way, the allelic differentiation Δ*D* was decomposed into differentiation among populations within a region (Δ*D*^1^), and differentiation among regions within an ecosystem (Δ*D*^2^). The details about the equations for diversity decomposition can be found in Gaggiotti *et al*, (2018).

As Shannon entropy avoids undue emphasis on either rare or common alleles (Sherwin *et al*. 2017), it is increasingly used in evolutionary biology and molecular ecology as a measure of genetic diversity and evolvability (Hampe, Schreiber & Krawczak 2003; Day 2015; Wagner 2017). Therefore, we also use summary statistics based on Shannon entropy (^1^*H*, ^*H*^*SS* hereafter) for comparison with diversity measures 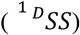. Shannon entropy per population (*H*^*P*^), per region (*H*^*R*^), and total Shannon entropy (*H*^*T*^) were calculated in line with the same hierarchies above. The additive decomposition of Shannon beta entropy (*H*_*β*_= *H*_*γ*_ - *H*_*α*_), was estimated at the population level 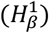 and regional level 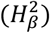 as well. Here, we also included Shannon differentiation (Δ*D*) to keep the number of statistics in ^*H*^*SS* the same with 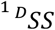.

In addition, we also calculated Mantel’s *r*, the correlation coefficient between genetic distance and geographical distance for *β*- measures 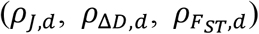, with distance measured in terms of the number of steps (edges) separating any two populations (vertices).

Each set of summary statistics includes the mean and standard deviation (*SD*). For each measure at the population level, we calculated the value for each population and the mean across populations. The total number of summary statistics for ^*Ar*^*SS* is 44, the same as for ^*He*^*SS*. The total number of summary statistics for 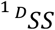 is 48 the same number as for ^*H*^*SS*. The description of summary statistics is shown in Table S1.

### Data analysis

The pipelines (R functions) to calculate the summary statistics are wrapped in the R package *HierDpart* (Qin 2019). We built 9 subsets of summary statistics ^*Ar*^*SS*, ^*H*^*SS*, ^*He*^*SS*, 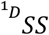, 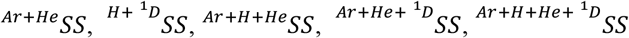, for the discriminatory power test.

### The power of summary statistics to discriminate among spatial scenarios

The power assessed by various machine learning methods may differ. Thus, to ensure that our tests are as comprehensive as possible, we employed three current state-of-the-art approaches to evaluate the power of the different subsets of summary statistics to discriminate among spatial structure scenarios: Kernel Local Fisher discriminant analysis (KLFDA; (Sugiyama 2007), conditional random forest classification (CRFC; (Strobl *et al*. 2007), and deep neural network (Ripley & Hjort 1996).

### Kernel Local Fisher Discriminant Analysis (KLFDA)

KLFDA is a recently proposed method for supervised dimensionality reduction based on local Fisher discriminant analysis (LFDA, Sugiyama 2006). As opposed to the standard Fisher discriminant analysis (LDA), LFDA can separate different classes (e.g. genetic clusters) while preserving the within-class structure (Sugiyama 2007); in other words, it allows for genetic sub-structuring within clusters. KLFDA represents an extension of LFDA that considers non-linear boundaries between classes through a nonlinear mapping of data points onto a reproducing kernel Hilbert space.

We carried out KLFDA on the 9 subsets of summary statistics. The Gaussian kernel was chosen for kernel transformation. Three key hyperparameters impact the accuracy of KLFDA, *d*, the number of reduced features for discriminant analysis, σ, the radius (the standard deviation) of the Gaussian kernel, and *knn*, the number of nearest neighbours. We first determined the appropriate number of reduced features ranging from 5 to 50 based on classification accuracy during training. We then did fine hyperparameter tuning on *σ* and *knn* via cross-validation with the best number of reduced features selected in the first step. *σ* value was tuned considering values between 0.001-10 (0.001, 0.005, 0.01, 0.05, 0.1, 0.5, 1, 5, 10) and *knn* was tuned between 5-50 (5, 10, 15, 20, 25, 30, 35, 40, 45, 50). Discriminatory power was evaluated by the classification accuracy (proportion of the overall correct discrimination) via leave-one-out cross-validation (Schaffer 1993; Kohavi 1995). Analyses were implemented using R package *lfda* (Tang & Li 2016; Tang & Li 2017).

### Conditional Random Forest Classification (CRFC)

We conducted the unbiased random forest classification based on conditional inference trees (*cforest*) that adopt the subsampling validation process with unbiased variable selection (bootstrap without replacement; (Strobl *et al*. 2007). To avoid overfitting in random forest classification, we optimized the key parameter (*mtry*) that governs the number of features that are randomly chosen to grow each tree from the bootstrapped data. We tuned the parameter *mtry* [mtry ϵ (1: *n*), *n* is the number of variables] via leave-one-out validation with 1000 trees for each subset of summary statistics. The parameter with the lowest average prediction error was chosen as the final model.

The standardized conditional importance of each variable, measured by the mean decrease in accuracy (MDA), was estimated from the optimum model based on bootstrapping without replacement according to (Strobl *et al*. 2008). Analyses were implemented using the R package “*caret*” (Kuhn 2015) calling *cforest* function from *party* package (Hothorn *et al*. 2010).

### Deep neural network

We conducted neural network (Baum 1988; Guarnieri, Piazza & Uncini 1999) classification using a 3 hidden layers perceptron (MLP) feedforward network with a weight decay to test the performance of the above subsets of summary statistics for spatial structure inference. The deep neural network training was carried out through a backpropagation with weighted decay optimization (a procedure to repeatedly adjust the weights to minimize the difference between true values and observed values) and a non-linear activation function (logistic) at the output layer. We first did a grid search on the parameter space via cross-validation to minimize the parameter range, then we tuned the parameters through dense parameter combinations via leave-one-out cross-validation. Finally, we tuned the number of neurons in each hidden layer using: layer1 = (1, 5, 10, 15); layer2= (0, 5, 10, 15); layer3= (0, 5, 10, 15), and the rate of decay using: decay = (0, 1e-5, 1e-4, 1e-3, 1e-2, 1e-1). Final model performance was evaluated by the model accuracy and Cohen’s Kappa coefficient (Cohen 1960). Models with the highest accuracy were chosen as the optimal model.

The (overall) importance of summary statistics is determined based on Garson’s algorithm (Garson 1991; Gevrey, Dimopoulos & Lek 2003), which uses combinations of the absolute values of the weights. We also used neural networks to assess the importance of summary statistics to identify a specific scenario. Deep neural network models were built using *caret* package (Kuhn 2008; Kuhn 2012).

### Evaluating discriminatory power of different sets of summary statistics

In terms of KLFDA, random forest classification, and neural network, we calculated the confusion matrix as well as overall performance statistics for each set of summary statistics. These model metrics are presented in Supplementary Material. Overall performance statistics included model accuracy and Kappa. All the performance statistics were estimated using the best model after leave-one-out cross-validation. The detailed description of these statistics can be found in (Kuhn & Johnson 2013).

We compared the performance of different sets of summary statistics in discriminating the five spatial scenarios using each of the above-mentioned methods separately to identify the best set of summary statistics.

## Results

### KLFDA inference

Table 2 presents the overall performance of different sets of summary statistics in discriminating five spatial scenarios using KLFDA. The best performing statistics set should have the highest accuracy and the largest Kappa value. Results indicated that 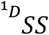 surpassed other sets of summary statistics at discriminating among scenarios and presented the highest discriminatory power. Though ^*H*^*SS* did slightly better than ^*Ar*^ *SS* and ^*He*^*SS*, it underperformed 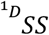 (Table 2). On the other hand, the set of summary statistics with the lowest discriminatory power corresponded to the most commonly used ^*He*^*SS* in population genetics (Table 2).

**Table 2.**
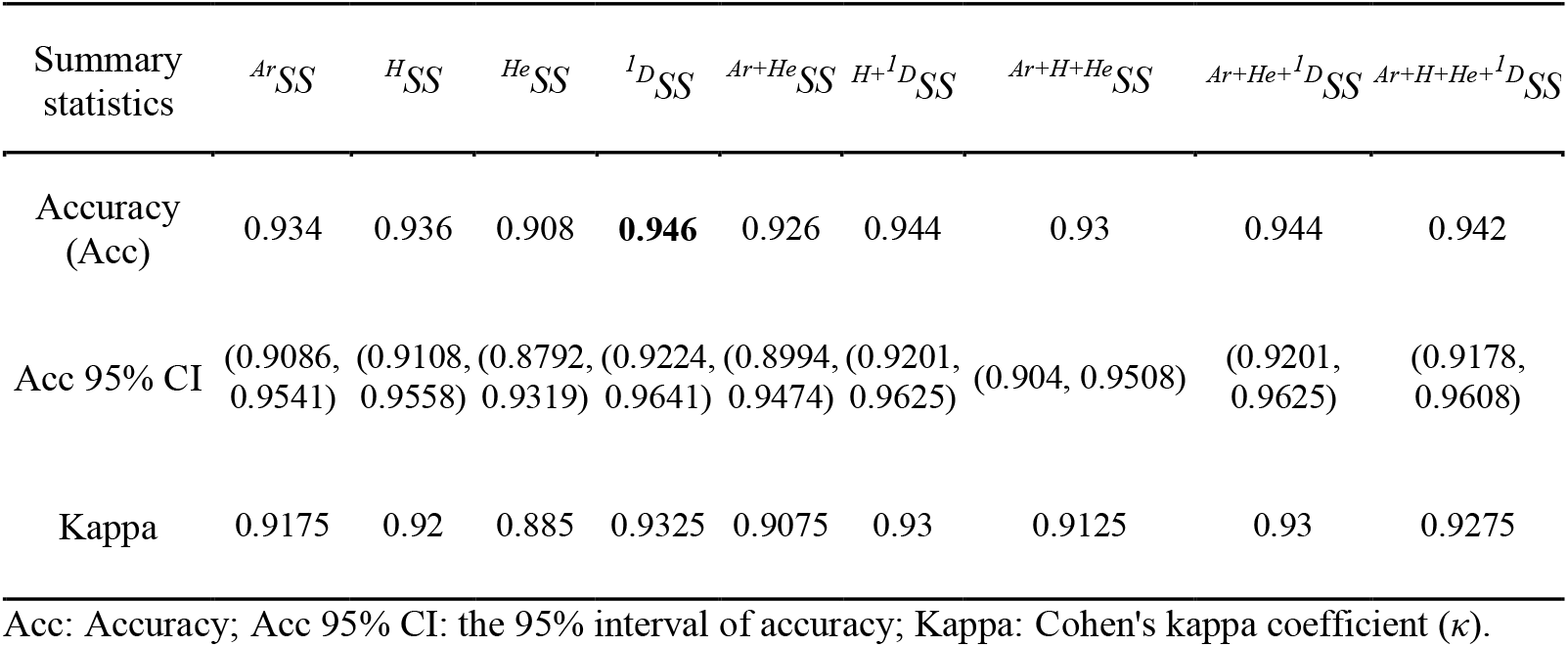
The overall performance of different sets of summary statistics in discriminating five spatial scenarios using KLFDA

Figure 1 presents results for the five scenarios based on the first two reduced features from KLFDA. Except 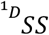, all other summary statistics, or the combination thereof, either failed to clearly distinguish between panmixia and the island model or failed to clearly distinguish between the standard stepping-stone model and hierarchical stepping-stone model (Fig. 1A-I). 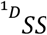 did a better job at discriminating among all of them (Fig. 1D).

**Fig. 1.**
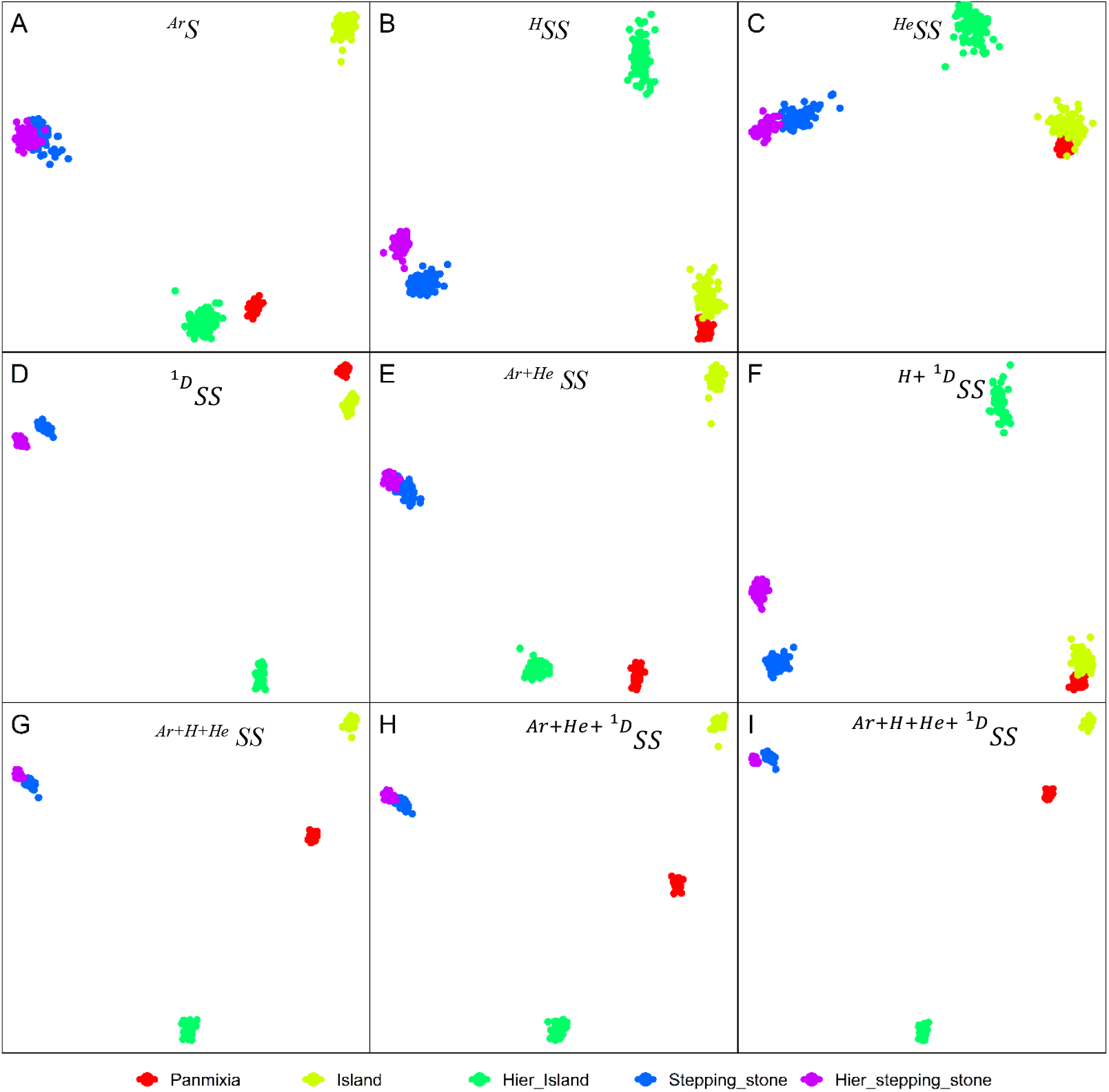
Projections of 5 spatial scenarios into two-dimensional subspaces using KLFDA based on different sets of summary statistics: 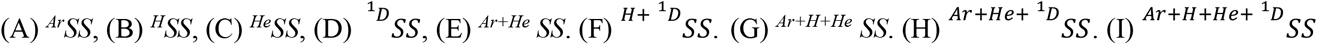. Each dot represents a simulated data set.

The confusion matrix supported these results (Table S2). Specifically, ^*Ar*^*SS* can correctly identify the island model, panmixia, and stepping-stone model (100%). But it did worse in identifying the hierarchical stepping-stone model (Fig. 1B, Table S2). ^*H*^*SS* did better at identifying hierarchical scenarios but performed less well in the case of the stepping-stone model (Fig. 1C, Table S2). 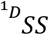 exhibited impressive performance across all scenarios with the exception of the hierarchical stepping-stone (Fig. 1D, Table S2). ^*He*^*SS* performed poorly in most scenarios with the exception of panmixia and hierarchical island scenarios (Fig. 1E, Table S2). Combinations of 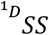 with other summary statistics showed similar results to those obtained with 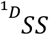 alone except when including ^*He*^*SS*, in which case discriminatory power was decreased (Table 2 & S2). In fact, combining ^*He*^*SS* with other summary statistics decreased the discriminatory power. Overall, the hierarchical stepping-stone scenario was the most difficult to identify correctly. 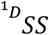 and ^*H*^*SS* did better at discriminating hierarchical stepping-stone model from other scenarios (Fig. 1, Table S2).

### Conditional Random Forest Classification

As is the case for KLFDA, among all the sets of summary statistics, ^*He*^*SS* had the lowest classification accuracy (Table 3). Slightly different from KLFDA results, 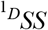 and ^*H*^*SS*, having the same discriminatory power, outclassed ^*Ar*^*SS* and ^*He*^*SS* in discriminating the five scenarios (Table 3). Note that conditional random forest didn’t show a power difference between 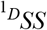 and ^*H*^*SS*, as well as between ^*Ar*+*H*+*He*^*SS*, 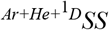, and 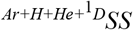(Table 3).

**Table 3.** The performance of different sets of summary statistics in discriminating five spatial scenarios using conditional random forest classification

| Summary statistics | ${}^{Ar}SS$ | ${}^HSS$ | ${}^{He}SS$ | ${}^1DSS$ | ${}^{Ar+He}SS$ | ${}^{H+{}^1D}SS$ | ${}^{Ar+H+He}SS$ | ${}^{Ar+He+{}^1D}SS$ | ${}^{Ar+H+He+{}^1D}SS$ |
| --- | --- | --- | --- | --- | --- | --- | --- | --- | --- |
| **Accuracy (Acc)** | 0.96 | 0.972 | 0.958 | 0.972 | 0.97 | 0.978 | **0.986** | **0.986** | **0.986** |
| **Acc 95% CI** | (0.9389, 0.9754) | (0.9535, 0.9846) | (0.9365, 0.9738) | (0.9535, 0.9846) | (0.951, 0.9831) | (0.961, 0.989) | (0.9714, 0.9944) | (0.9714, 0.9944) | (0.9714, 0.9944) |
| **Kappa** | 0.95 | 0.965 | 0.9475 | 0.965 | 0.9625 | 0.9725 | 0.9825 | 0.9825 | 0.9825 |

Acc: Accuracy; Acc 95% CI: the 95% interval of accuracy; Kappa: Cohen's kappa coefficient ( $\kappa$ ).

Compared to KLFDA, the discriminatory power of all sets of summary statistics to discriminate spatial scenarios increased when using conditional random forest (Table 3). Moreover, as opposed to KLFDA results, combining different sets of summary statistics led to an increase in discriminatory power (Table 3). The most difficult scenario to identify is the stepping-stone model. However, consistent with KLFDA results, ^*Ar*^*SS* showed the worse performance to distinguish the stepping-stone scenario than ^*H*^*SS*, 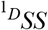 and ^*He*^*SS* (Table S3). ^*He*^*SS* did a worse job at identifying hierarchical stepping-stone model compared to ^*Ar*^*SS*, ^*H*^*SS*, and 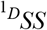 (Table S3).

A particular advantage of random forest classification is that it allows us to rank individual summary statistics in terms of their discriminating power. Figure 2 presents the top 30 ranked summary statistics among the total 178 summary statistics including ^*Ar*^*SS*, ^*H*^*SS*, ^*He*^*SS*, and 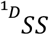. The best performing statistics in discriminating the spatial scenarios were the *β*-measures and their Mantel statistics (*ρ*). Among all the summary statistics, *SD*(*ρ*_*J,d*_) (belonging to ^*Ar*^*SS*) and *ρ*_*ΔD,d*_ (belonging to 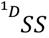 were the two most important statistics contributing to the ability to discriminate among all spatial scenarios (Fig. 2). Four out of the top-ten ranked statistics, *SD*(*ρ*_*j,d*_), *ρ*_*j,d*_, 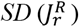, and 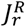, accounting for first, third, fourth, and seventh best-performing statistics respectively, belong to ^*Ar*^*SS*. Three out of the top-ten ranked statistics *ρ*_*ΔD,d*_, *SD*(*ρ*_*ΔD,d*_), and *SD*(Δ*D*^2^) from 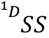 and ^*H*^*SS*, accounted for the second, fifth, and eighth most important statistics respectively. The last three statistics out of the top-ten, *ρ*_*Fst,d*_, 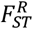, and 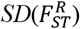, which belong to ^*He*^*SS*, had relatively low importance when compared to ^*Ar*^*SS* and 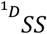 and ranked as the sixth, ninth, and tenth best-performing statistics respectively (Fig. 2).

**Fig. 2.**
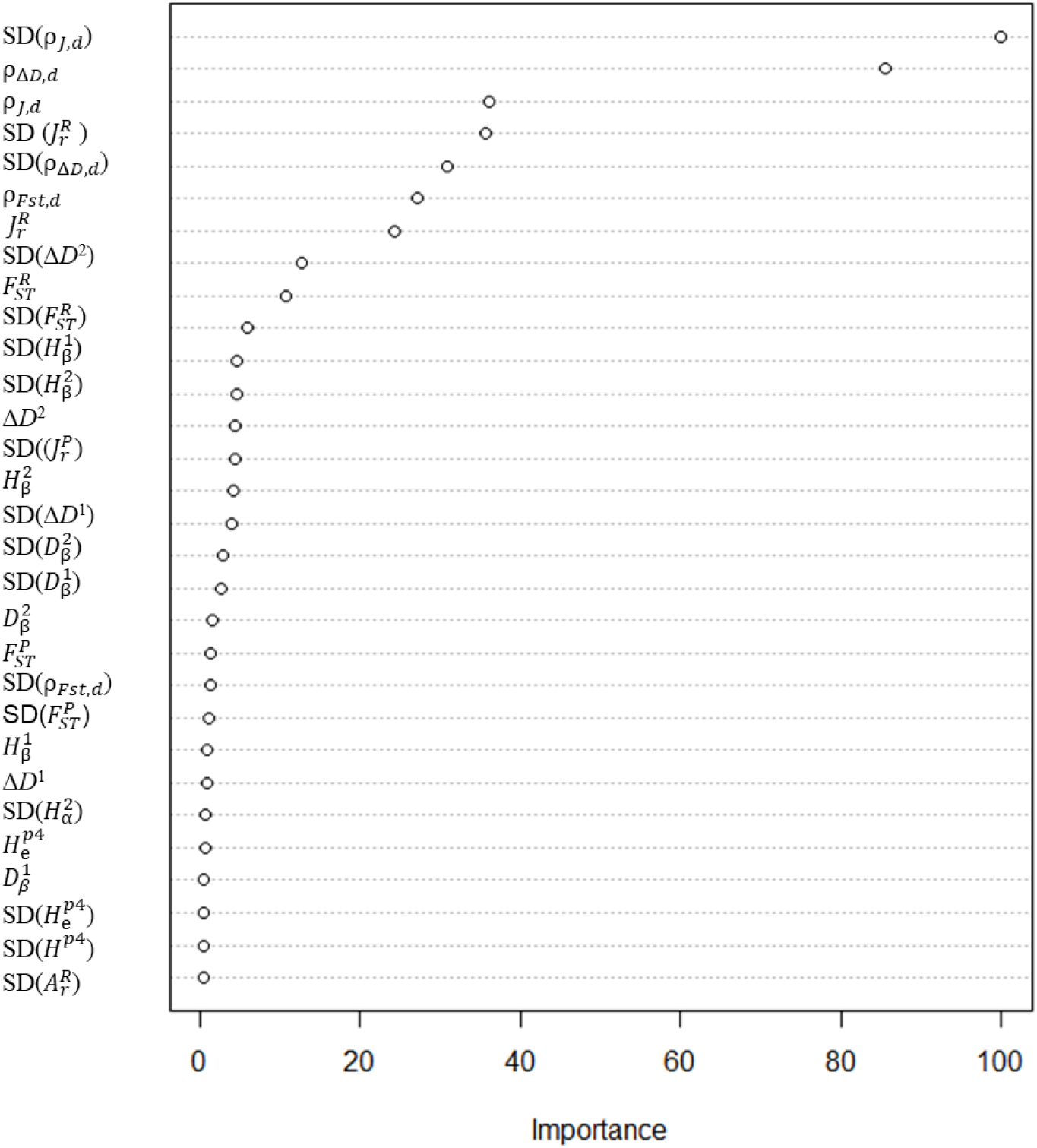
Ranked conditional variable importance estimated by conditional random forest classification. Results are shown only for the top 30 most important summary statistics among the 178 summary statistics. Statistics abbreviations are given in Table S1.

### Deep neural network

The deep neural network analysis produced results similar to those of the two previous methods. Generally, the summary statistics can be categorized into four discriminatory sets based on discriminatory power. Again, 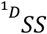, the most powerful summary statistics, along with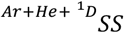, outclassed other sets of summary statistics (Table 4). ^*Ar*^*SS*, ^*H*^*SS* and ^*Ar*+*H*+*He*^*SS*, comprised the second most discriminatory sets of summary statistics, with their discriminant accuracy being only slightly lower than 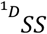 (0.988, Table 4). The third most discriminatory sets of summary statistics were ^*Ar*+*He*^*SS*, ^*H*+*D*^*SS*, and 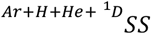. Finally, the least discriminatory set of summary statistics was ^*He*^*SS* (Table 4). As it was the case with KLFDA, neural network results indicated that combining different sets of summary statistics (increasing the number of summary statistics) did not increase discriminatory power (Table 4).

**Table 4.**
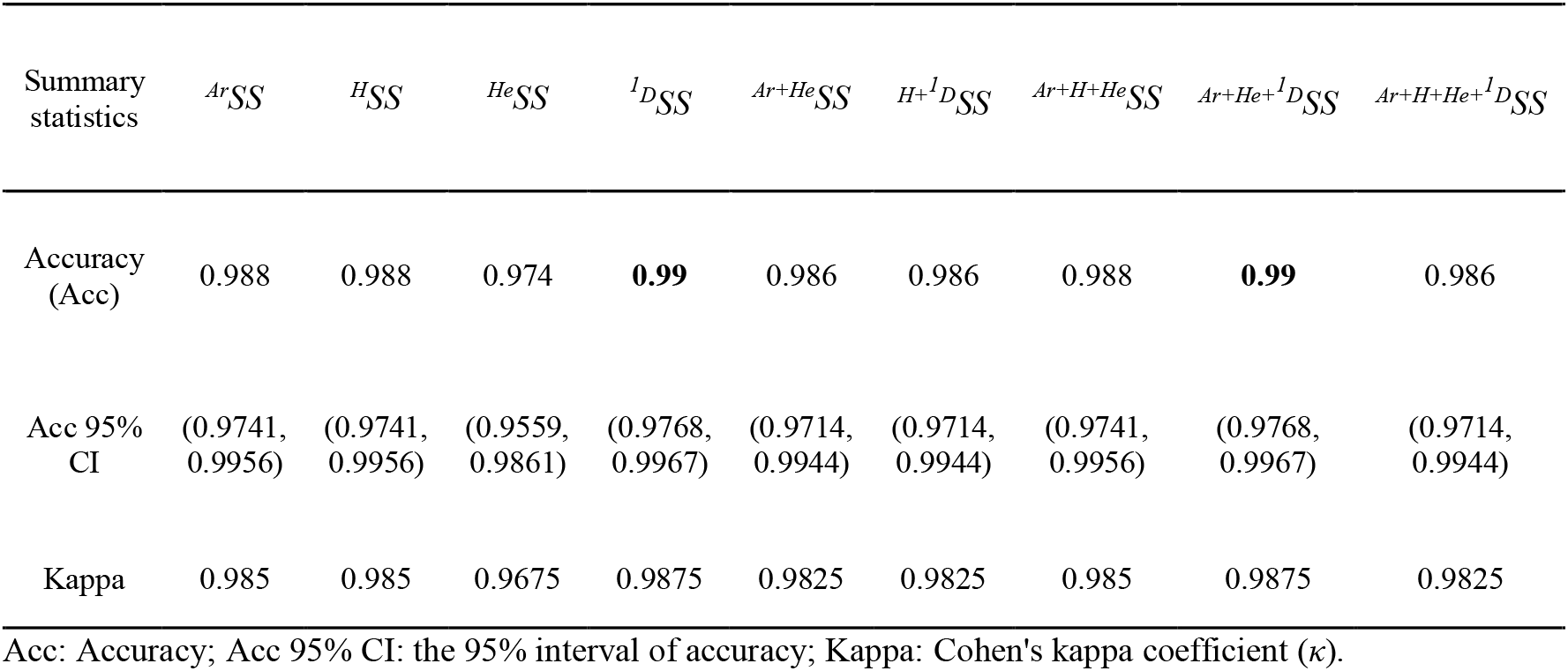
Performance of different sets of summary statistics in discriminating five spatial scenarios using deep neural network

The discriminatory power of all sets of summary statistics using the neural network was higher than that of KLFDA and CRFC (Tables 2-4). This indicates that the neural network performed better than the two other ML methods. Unlike KLFDA and CRFC, the deep neural network did better at discriminating between panmixia and island model, with most sets of summary statistics (except ^*He*^*SS*) 100 % successfully discriminating between these two scenarios (Table S4). 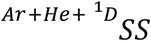 and ^*Ar*+*H*+*He*^*SS* did a better job (0.2% error rate) in differentiating the stepping-stone model and hierarchical stepping-stone model compared to other sets of summary statistics (Table S4).

Figure 3 presents the variable importance of the top 30 ranked summary statistics among the total 178 summary statistics according to their discriminatory power estimated from the deep neural network. ^*H*^*SS*, 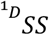, ^*Ar*^*SS* and ^*He*^*SS* accounted for 11/30 (5 overlapped statistics with 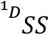), 10/30, 8/30, and 6/30 of the top-30 ranked summary statistics respectively (Fig. 3). The first three most informative summary statistics were 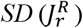 and *ρ*_*ΔD,d*_. They contributed equally toward the ability to discriminate among all spatial scenarios (importance values are all 100, Figs. 3 & S1). Similar to CRFC results, among the top 10 most informative statistics, the first (*ρ*_*J,d*_), second 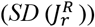, and the seventh (*SD*(*ρ*_*J,d*_)) most important statistics belong to ^*Ar*^*SS. ρ*_*ΔD,d*_, *SD*(*ρ*_*ΔD,d*_)), *SD*(Δ*D*^2^), and 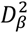, which were the third, fourth, sixth, and tenth best-performing summary statistics respectively, belong to 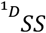. Only two out of ten best-performing statistics, *ρ*_*Fst,d*_ and 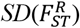, ranking as the fifth and the eighth-most important summary statistic respectively, belong to ^*He*^*SS* (Fig. 3).

**Fig. 3.**
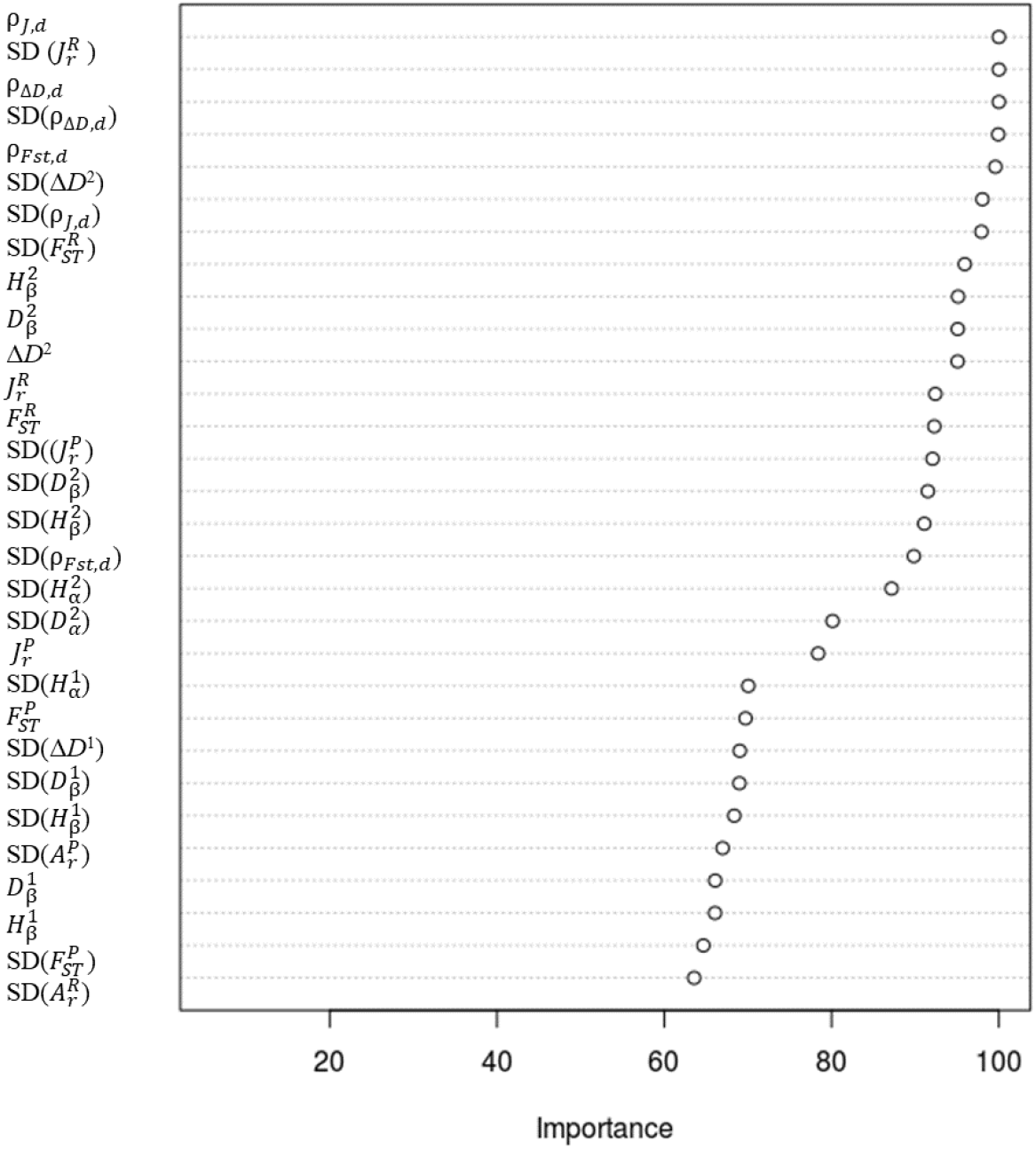
Variable importance estimated using the deep neural network. Results are shown only for the top 30 ranked summary statistics among the 178 summary statistics. Statistics abbreviations are given in Table S1.

Figure S1 presents the scenario-specific variable importance ranked in accordance with their overall importance (c.f., Fig. 3). The 16 top summary statistics contributed almost equally to panmixia, stepping-stone model, hierarchical stepping-stone model and hierarchical island model (Fig. S1). On the other hand, only the top five statistics, 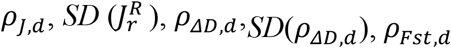, contributed most to the power of discriminating the island model from other models (Fig. S1). Besides the top 16 most important statistics, 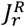 and 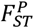 also contributed substantially to the power of discriminating stepping-stone and hierarchical stepping-stone models (Fig. S1).

## Discussion

In this study, we performed a comprehensive assessment of the discriminatory power of 9 sets of summary statistics, comprised of ^*Ar*^*SS*, ^*H*^*SS*, ^*He*^*SS* and 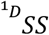. Since different methods to estimate discriminatory power may lead to different results, we employed three up-to-date machine learning methods to compare the power of the different sets of summary statistics. All results led to the same conclusion that 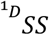 outperformed the other sets of summary statistics in the discrimination of spatial-structure scenarios. Though, 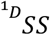, was overall the best set of diversity measures, without undue emphasising on rare or common entities, ^*Ar*^*SS* and ^*He*^*SS* also provided complementary information that 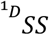 did not capture.

Jaccard dissimilarity (*J*) and its Mantel’ *r* ranked as the top summary statistics among all the summary statistics for differentiating spatial scenarios, followed by Δ*D* and then *F*_*ST*_ as well as their Mantel’s *r*. In addition, we found that combining sets of summary statistics did not necessarily increase discriminatory power (e.g., KLFDA and neural network models in Tables 2 & 4). Therefore, a more efficient strategy would be combining the most informative summary statistics in each set depending on the alternative spatial scenarios that could apply to each dataset based on existing information.

During the past 20 years, evolutionary biologists and population geneticists have been using diversity metrics as the summary statistics to make inference on the evolutionary and demographic histories of populations via approximate Bayesian computation (ABC). Information theory offers a spectrum of summary statistics that can be used with ABC. However, the choice of summary statistics in population genetics has focused on the ^*He*^*SS* family (i.e., heterozygosity, *He*, and fixation, *Fst*). The use of ^*He*^*SS* up-weight the signal provided by common alleles while down-weighting rare alleles, thus it may miss important information under scenarios that involve bottlenecks or founder events. To avoid this problem, it is common to combine ^*Ar*^*SS* and ^*He*^*SS*, however, our results indicate that the same or more discriminatory power could be obtained using only the 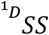 set. These results provide further support for the idea that simply increasing the number of summary statistics without considering their individual discriminatory power may decrease the inference accuracy.

Our systematic assessment of the power of these summary statistics showed that, ^*He*^*SS*, the most commonly used set of summary statistics in population genetics performed worst in the discrimination of typical spatial-structure scenarios tested by three different classification approaches. *J*, Δ*D*, and *F*_*ST*_ are *β*-diversity measures evaluating the extent of genetic differentiation between populations, with Δ*D* and *F*_*ST*_ being estimated based on allele frequency, and *J* being estimated based on allele presence/absence data. Generally, genetic differentiation is usually estimated using *F*_*ST*_ (Wright 1949) and its variants (*G*_*ST*_ (Nei 1973)) calculated from heterozygosity (*He*) while *J* and Δ*D*, which are more informative according to our results, are rarely used as statistics to measure population genetic inference.

Our results indicate that ^*Ar*^*SS* contributed better to differentiate between panmixia and the other scenarios. 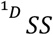 on the other hand, exhibits high accuracy in differentiating all scenarios, especially being good at discriminating between stepping-stone and hierarchical stepping-stone models. Therefore, ^*Ar*^*SS* seems useful for detecting the scenarios that depart from panmixia, and 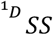 may be helpful to differentiate between more complex spatial scenarios. On the other hand, we did not observe advantageous properties in ^*He*^*SS* in detecting the spatial structuring signals under the five spatial scenarios considered. Though *ΔD* showed high power of detecting the signal of the spatial structure changes, there is still a lack of knowledge about the relationship between *ΔD* and demographic parameters, as well as *ΔD*’s response to selection.

For a long time, important guidelines for species and genetic diversity conservation have been made using the richness and Simpson index in terms of species diversity (Scott *et al*. 1987; Jost *et al*. 2010), and heterozygosity (derived from *F*-statistics framework) in terms of genetic diversity (Aitken, Luikart & Allendorf 2012). The results of this study suggest that summary statistics based on Hill’s numbers are promising tools for detecting diversity changes in biological conservation studies.

In summary, diversity of order *q* =1 (^1^ *D*) and Shannon differentiation offer a unified approach integrating diversity across all levels of biological organizations. Our results suggest that 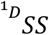 would perform well for the purpose of inference of population structure using inferential frameworks such as approximate Bayesian computation (ABC). It is clear that no single set of diversity measures can capture all the information contained in raw population genetics datasets and our study suggest that the type of summary statistic we may want to use depends on the specific question being asked.

Finally, we found different machine learning methods showed different performance to distinguish spatial structure scenarios. KLFDA gave the lowest discriminant accuracy while the deep neural network gave the highest discriminant accuracy among the three classification methods (Tables 2-4). In contrast, conditional random forest did not show the difference between the power of ^*H*^*SS* and 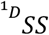 as well as other combinations of summary statistics (Table 3). The conditional random forest also showed lower power to identify the importance of summary statistics compared to neural networks (Figs. 2-3). The deep neural network showed more advantages than KLFDA and conditional random forest in this study, which provides additional support to recent assertions that machine learning methods represent promising tools to carry out inference in ecology and evolution (Schrider & Kern 2018).

## Data and code availability

The input files and scripts for generating simulation as well as the analyses of summary statistics are available at https://github.com/xinghuq/SS_performance.

## Acknowledgments

XHQ was supported by China Scholarship Council.

## Author contribution

XQ and OEG designed the study. XQ carried out the analyses and interpreted results with the input from OEG. XQ wrote the manuscript with the input of OEG. Both authors contributed to editing and revising the manuscript.

