## Supplementary Materials for "Information-based summary statistics for spatial genetic structure inference"

**Supplementary tables and figures**

**Table S1**. List of summary statistics

| **Hierarchical levels** | **Acronym** | **Traditional summary statistics** | **#statistics** | **Acronym** | **New summary statistics (based on order *q*=1)** | **#statistics** |
| --- | --- | --- | --- | --- | --- | --- |
|  | $A_{r}^{pi}$ | Allelic richness of each population | 16 | $D^{pi}$ | Allelic diversity of each population | 16 |
|  | *SD* ($A_{r}^{pi}$) | Standard deviation of allelic richness of each population | 16 | ${SD(D}^{pi})$ | Standard deviation of allelic diversity of each population | 16 |
|  | $A_{r}^{P}$ | Average allelic richness per population | 1 | $D_{\alpha}^{1}$ | Average allelic diversity per population | 1 |
|  | *SD* ($A_{r}^{P}$) | Standard deviation of average allelic richness per population | 1 | *SD* ($D_{\alpha}^{1}$ ） | Standard deviation of average allelic diversity per population | 1 |
| Population | $H_{e}^{pi}$ | Expected heterozygosity of each population | 16 | $D_{\beta}^{1}$ | Equivalent number of populations | 1 |
|  | $SD\left( H_{e}^{pi} \right)$ | Standard deviation of expected heterozygosity of each population | 16 | *SD* ($D_{\beta}^{1}$ ) | Standard deviation of equivalent number of populations | 1 |
|  | $H_{e}^{P}$ | Average expected heterozygosity per population | 1 | $H^{pi}$ | Shannon entropy of each population | 16 |
|  | $SD\left( H_{e}^{P} \right)$ | Standard deviation of average expected heterozygosity per population | 1 | *SD* ($H^{pi}$) | Standard deviation of Shannon entropy of each population | 16 |
|  |  |  |  | $H^{P}$ | Average Shannon entropy per populations | 1 |
|  |  |  |  | *SD* ($H^{P}$) | Standard deviation of average Shannon entropy per populations | 1 |
|  |  |  |  | $H_{\beta}^{1}$ | Beta Shannon entropy between pops within region | 1 |
|  |  |  |  | *SD* ( $H_{\beta}^{1})$ | Standard deviation of beta Shannon entropy between pops within region | 1 |
|  |  |  |  | Genetic structure (*β*-diversity) |  |  |
| Between population within region | $J_{r}^{P}$ | *Jaccard* dissimilarity between populations | 1 | *ΔD*^1^ | Allelic differentiation between populations | 1 |
|  | *SD* ($J_{r}^{P}$) | Standard deviation of *Jaccard* dissimilarity between populations | 1 | *SD* (*ΔD*^1^) | Stand deviation of allelic differentiation between populations | 1 |
|  | $F_{ST}^{P}$ | Genetic fixation between populations | 1 |  |  |  |
|  | *SD* ($F_{ST}^{P}$) | Standard deviation of genetic fixation between populations | 1 |  |  |  |
| Correlation between differentiation and distance | $\rho_{J,d}$ | Correlation between Jaccard dissimilarity and geographic distance | 1 | $\rho_{\Delta D,d}$ | Correlation between Δ*D* and geographic distance | 1 |
|  | *SD* ($\rho_{J,d}$) | Standard deviation of correlation between Jaccard dissimilarit*y* and geographic distance | 1 | *SD* ($\rho_{\Delta D,d}$) | Standard deviation of correlation between Δ*D* and geographic distance | 1 |
|  | $\rho_{Fst,d}$ | Correlation between **F***_ST_* and geographic distance | 1 |  |  |  |
|  | *SD* ($\rho_{Fst,d}$) | Standard deviation of correlation between **F***_ST_* and geographic distance | 1 |  |  |  |
| Region | $A_{r}^{R}$ | Average allelic richness per region | 1 | $D_{\alpha}^{2}$ | Average allelic diversity per region | 1 |
|  | *SD* ($A_{r}^{R}$) | Standard deviation of average allelic richness per region | 1 | *SD* ($D_{\alpha}^{2}$) | Standard deviation of average allelic diversity per region | 1 |
|  | $H_{e}^{R}$ | Average expected heterozygosity per region | 1 | $D_{\beta}^{2}$ | Equivalent number of regions | 1 |
|  | *SD* ($H_{e}^{R}$) | Standard deviation of average expected heterozygosity per region | 1 | *SD* ($D_{\beta}^{2}$) | Standard deviation of equivalent number of regions | 1 |
|  |  |  |  | $H^{R}$ | Average Shannon entropy per region | 1 |
|  |  |  |  | *SD* ($H^{R}$) | Standard deviation of average Shannon entropy per region | 1 |
|  |  |  |  | $H_{\beta}^{2}$ | Beta Shannon entropy among regions | 1 |
|  |  |  |  | *SD* ($H_{\beta}^{2})$ | Standard deviation of beta Shannon entropy among regions | 1 |
|  |  |  |  | Genetic structure (*β*-diversity) |  |  |
| Between regions within ecosystem | $J_{r}^{R}$ | Jaccard dissimilarity between regions within ecosystem |  | *ΔD*^2^ | Allelic differentiation between regions within ecosystem | 1 |
|  | *SD* ($J_{r}^{R}$) | Standard deviation of Jaccard dissimilarity between regions within ecosystem |  | *SD* (*ΔD*^2^) | Stand deviation of allelic differentiation between regions within ecosystem | 1 |
|  | $F_{ST}^{R}$ | Genetic fixation between regions within ecosystem | 1 |  |  |  |
|  | *SD* ($F_{ST}^{R}$) | Standard deviation of genetic fixation between regions within ecosystem | 1 |  |  |  |
| Ecosystem | $H_{e}^{T}$ | Total expected heterozygosity in the ecosystem | 1 | *D_γ_* | Total allelic diversity in the ecosystem | 1 |
|  | *SD* ($H_{e}^{T}$) | Standard deviation of total expected heterozygosity | 1 | *SD* (*D_γ_*) | Standard deviation of total allelic diversity | 1 |
|  | $A_{r}^{T}$ | Total allelic richness in the ecosystem |  | *H^T^* | Total Shannon entropy in the ecosystem | 1 |
|  | *SD* ($A_{r}^{T}$) | Standard deviation of total allelic richness in ecosystem |  | *SD* (*H^T^*) | Standard deviation of total Shannon entropy in the ecosystem | 1 |

**Table S2**. Confusion matrix of summary statistics tested by KLFDA

|  |  |  | *^Ar^SS* |  |  |
| --- | --- | --- | --- | --- | --- |
|  | Hier_Island | Hier_stepping stone | Island | Panmixia | Stepping_stone |
| Hier_Island | 97 | 1 | 0 | 0 | 0 |
| Hier_steppingstone | 1 | 70 | 0 | 0 | 0 |
| Island | 2 | 0 | 100 | 0 | 0 |
| Panmixia | 0 | 0 | 0 | 100 | 0 |
| Stepping_stone | 0 | 29 | 0 | 0 | 100 |
|  |  |  | *^H^SS* |  |  |
|  | Hier_Island | Hier_stepping stone | Island | Panmixia | Stepping_stone |
| Hier_Island | 99 | 0 | 4 | 0 | 0 |
| Hier_steppingstone | 0 | 90 | 2 | 0 | 15 |
| Island | 1 | 1 | 94 | 0 | 0 |
| Panmixia | 0 | 0 | 0 | 100 | 0 |
| Stepping_stone | 0 | 9 | 0 | 0 | 85 |
|  |  |  | *^He^SS* |  |  |
|  | Hier_Island | Hier_stepping stone | Island | Panmixia | Stepping_stone |
| Hier_Island | 97 | 3 | 7 | 0 | 0 |
| Hier_steppingstone | 2 | 84 | 1 | 0 | 16 |
| Island | 0 | 0 | 89 | 0 | 0 |
| Panmixia | 0 | 0 | 3 | 100 | 0 |
| Stepping_stone | 1 | 13 | 0 | 0 | 84 |
|  |  |  | ${}^{{}^{1}D}{SS}$ |  |  |
|  | Hier_Island | Hier_stepping stone | Island | Panmixia | Stepping_stone |
| Hier_Island | 96 | 0 | 0 | 0 | 0 |
| Hier_steppingstone | 1 | 82 | 0 | 0 | 1 |
| Island | 3 | 0 | 96 | 0 | 0 |
| Panmixia | 0 | 0 | 4 | 100 | 0 |
| Stepping_stone | 0 | 18 | 0 | 0 | 99 |
|  |  |  | *^Ar+He^ SS* |  |  |
|  | Hier_Island | Hier_stepping stone | Island | Panmixia | Stepping_stone |
| Hier_Island | 97 | 0 | 0 | 0 | 0 |
| Hier_steppingstone | 0 | 69 | 0 | 0 | 0 |
| Island | 3 | 0 | 97 | 0 | 0 |
| Panmixia | 0 | 0 | 3 | 100 | 0 |
| Stepping_stone | 0 | 31 | 0 | 0 | 100 |
|  |  |  | ${}^{H+{}^{1}D}{SS}$ |  |  |
|  | Hier_Island | Hier_stepping stone | Island | Panmixia | Stepping_stone |
| Hier_Island | 94 | 0 | 0 | 0 | 0 |
| Hier_steppingstone | 3 | 81 | 0 | 0 | 1 |
| Island | 3 | 0 | 98 | 0 | 0 |
| Panmixia | 0 | 0 | 2 | 100 | 0 |
| Stepping_stone | 0 | 19 | 0 | 0 | 99 |
|  |  |  | ${}^{Ar+H+He}{SS}$ |  |  |
|  | Hier_Island | Hier_stepping stone | Island | Panmixia | Stepping_stone |
| Hier_Island | 97 | 0 | 0 | 0 | 0 |
| Hier_steppingstone | 0 | 70 | 0 | 0 | 0 |
| Island | 3 | 0 | 98 | 0 | 0 |
| Panmixia | 0 | 0 | 2 | 100 | 0 |
| Stepping_stone | 0 | 30 | 0 | 0 | 100 |
|  |  |  | ${}^{Ar+He+{}^{1}D}{SS}$ |  |  |
|  | Hier_Island | Hier_stepping stone | Island | Panmixia | Stepping_stone |
| Hier_Island | 97 | 0 | 0 | 0 | 0 |
| Hier_steppingstone | 0 | 77 | 0 | 0 | 0 |
| Island | 3 | 0 | 98 | 0 | 0 |
| Panmixia | 0 | 0 | 2 | 100 | 0 |
| Stepping_stone | 0 | 23 | 0 | 0 | 100 |
|  |  |  | ${}^{Ar+H+He+{}^{1}D}{SS}$ |  |  |
|  | Hier_Island | Hier_stepping stone | Island | Panmixia | Stepping_stone |
| Hier_Island | 97 | 0 | 0 | 0 | 0 |
| Hier_steppingstone | 0 | 76 | 0 | 0 | 0 |
| Island | 3 | 0 | 98 | 0 | 0 |
| Panmixia | 0 | 0 | 2 | 100 | 0 |
| Stepping_stone | 0 | 24 | 0 | 0 | 100 |

**Table S3.** Confusion matrix of summary statistics from random forest classification (cforest)

|  |  |  | *^Ar^SS* |  |  |
| --- | --- | --- | --- | --- | --- |
|  | Hier_Island | Hier_stepping stone | Island | Panmixia | Stepping_stone |
| Hier_Island | 100 | 0 | 1 | 0 | 0 |
| Hier_steppingstone | 0 | 99 | 0 | 0 | 18 |
| Island | 0 | 0 | 99 | 0 | 0 |
| Panmixia | 0 | 0 | 0 | 100 | 0 |
| Stepping_stone | 0 | 1 | 0 | 0 | 82 |
|  |  |  | *^H^SS* |  |  |
|  | Hier_Island | Hier_stepping stone | Island | Panmixia | Stepping_stone |
| Hier_Island | 99 | 0 | 0 | 0 | 0 |
| Hier_steppingstone | 0 | 99 | 0 | 0 | 10 |
| Island | 1 | 0 | 98 | 0 | 0 |
| Panmixia | 0 | 0 | 2 | 100 | 0 |
| Stepping_stone | 0 | 1 | 0 | 0 | 90 |
|  |  |  | *^He^SS* |  |  |
|  | Hier_Island | Hier_stepping stone | Island | Panmixia | Stepping_stone |
| Hier_Island | 98 | 1 | 2 | 0 | 0 |
| Hier_steppingstone | 1 | 94 | 0 | 0 | 9 |
| Island | 1 | 0 | 97 | 1 | 0 |
| Panmixia | 0 | 0 | 1 | 99 | 0 |
| Stepping_stone | 0 | 5 | 0 | 0 | 91 |
|  |  |  | ${}^{{}^{1}D}{SS}$ |  |  |
|  | Hier_Island | Hier_stepping stone | Island | Panmixia | Stepping_stone |
| Hier_Island | 100 | 0 | 1 | 0 | 0 |
| Hier_steppingstone | 0 | 99 | 0 | 0 | 10 |
| Island | 0 | 0 | 97 | 0 | 0 |
| Panmixia | 0 | 0 | 2 | 100 | 0 |
| Stepping_stone | 0 | 1 | 0 | 0 | 90 |
|  |  |  | *^Ar+He^ SS* |  |  |
|  | Hier_Island | Hier_stepping stone | Island | Panmixia | Stepping_stone |
| Hier_Island | 99 | 0 | 2 | 0 | 0 |
| Hier_steppingstone | 0 | 99 | 0 | 0 | 11 |
| Island | 0 | 0 | 98 | 0 | 0 |
| Panmixia | 0 | 0 | 0 | 100 | 0 |
| Stepping_stone | 1 | 1 | 0 | 0 | 89 |
|  |  |  | ${}^{H+{}^{1}D}{SS}$ |  |  |
|  | Hier_Island | Hier_stepping stone | Island | Panmixia | Stepping_stone |
| Hier_Island | 100 | 0 | 0 | 0 | 0 |
| Hier_steppingstone | 0 | 99 | 0 | 0 | 8 |
| Island | 0 | 0 | 98 | 0 | 0 |
| Panmixia | 0 | 0 | 2 | 100 | 0 |
| Stepping_stone | 0 | 1 | 0 | 0 | 92 |
|  |  |  | *^Ar+H+He^ SS* |  |  |
|  | Hier_Island | Hier_stepping stone | Island | Panmixia | Stepping_stone |
| Hier_Island | 100 | 0 | 0 | 0 | 0 |
| Hier_steppingstone | 0 | 99 | 0 | 0 | 6 |
| Island | 0 | 0 | 100 | 0 | 0 |
| Panmixia | 0 | 0 | 0 | 100 | 0 |
| Stepping_stone | 0 | 1 | 0 | 0 | 94 |
|  |  |  | ${}^{Ar+He+{}^{1}D}{SS}$ |  |  |
|  | Hier_Island | Hier_stepping stone | Island | Panmixia | Stepping_stone |
| Hier_Island | 100 | 0 | 0 | 0 | 0 |
| Hier_steppingstone | 0 | 99 | 0 | 0 | 6 |
| Island | 0 | 0 | 100 | 0 | 0 |
| Panmixia | 0 | 0 | 0 | 100 | 0 |
| Stepping_stone | 0 | 1 | 0 | 0 | 94 |
|  |  |  | ${}^{Ar+H+He+{}^{1}D}{SS}$ |  |  |
|  | Hier_Island | Hier_stepping stone | Island | Panmixia | Stepping_stone |
| Hier_Island | 100 | 0 | 0 | 0 | 0 |
| Hier_steppingstone | 0 | 99 | 0 | 0 | 6 |
| Island | 0 | 0 | 100 | 0 | 0 |
| Panmixia | 0 | 0 | 0 | 100 | 0 |
| Stepping_stone | 0 | 1 | 0 | 0 | 94 |

**Table S4.** Confusion matrix of summary statistics tested by MLP neural network

|  |  | *^Ar^ SS* | |  |  |
| --- | --- | --- | --- | --- | --- |
|  | Hier_Island | Hier_stepping stone | Island | Panmixia | Stepping_stone |
| Hier_Island | 100 | 0 | 0 | 0 | 0 |
| Hier_steppingstone | 0 | 96 | 0 | 0 | 2 |
| Island | 0 | 0 | 100 | 0 | 0 |
| Panmixia | 0 | 0 | 0 | 100 | 0 |
| Stepping_stone | 0 | 4 | 0 | 0 | 98 |
| *^H^SS* | | | | | |
|  | Hier_Island | Hier_stepping stone | Island | Panmixia | Stepping_stone |
| Hier_Island | 100 | 0 | 0 | 0 | 0 |
| Hier_steppingstone | 0 | 97 | 0 | 0 | 3 |
| Island | 0 | 0 | 100 | 0 | 0 |
| Panmixia | 0 | 0 | 0 | 100 | 0 |
| Stepping_stone | 0 | 3 | 0 | 0 | 97 |
| *^He^SS* | | | | | |
|  | Hier_Island | Hier_stepping stone | Island | Panmixia | Stepping_stone |
| Hier_Island | 96 | 1 | 0 | 0 | 0 |
| Hier_steppingstone | 0 | 96 | 0 | 0 | 4 |
| Island | 4 | 0 | 99 | 0 | 0 |
| Panmixia | 0 | 0 | 1 | 100 | 0 |
| Stepping_stone | 0 | 3 | 0 | 0 | 96 |
| ${}^{{}^{1}D}{SS}$ | | | | | |
|  | Hier_Island | Hier_stepping stone | Island | Panmixia | Stepping_stone |
| Hier_Island | 100 | 0 | 0 | 0 | 0 |
| Hier_steppingstone | 0 | 97 | 0 | 0 | 2 |
| Island | 0 | 0 | 100 | 0 | 0 |
| Panmixia | 0 | 0 | 0 | 100 | 0 |
| Stepping_stone | 0 | 3 | 0 | 0 | 98 |
| *^Ar+He^ SS* | | | | | |
|  | Hier_Island | Hier_stepping stone | Island | Panmixia | Stepping_stone |
| Hier_Island | 100 | 0 | 0 | 0 | 0 |
| Hier_steppingstone | 0 | 96 | 0 | 0 | 3 |
| Island | 0 | 0 | 100 | 0 | 0 |
| Panmixia | 0 | 0 | 0 | 100 | 0 |
| Stepping_stone | 0 | 4 | 0 | 0 | 97 |
| ${}^{H+{}^{1}D}{SS}$ | | | | | |
|  | Hier_Island | Hier_stepping stone | Island | Panmixia | Stepping_stone |
| Hier_Island | 98 | 0 | 0 | 0 | 0 |
| Hier_steppingstone | 0 | 97 | 0 | 0 | 2 |
| Island | 2 | 0 | 100 | 0 | 0 |
| Panmixia | 0 | 0 | 0 | 100 | 0 |
| Stepping_stone | 0 | 3 | 0 | 0 | 98 |
| *^Ar+H+He^ SS* | | | | | |
|  | Hier_Island | Hier_stepping stone | Island | Panmixia | Stepping_stone |
| Hier_Island | 98 | 0 | 0 | 0 | 0 |
| Hier_steppingstone | 0 | 98 | 0 | 0 | 2 |
| Island | 2 | 0 | 100 | 0 | 0 |
| Panmixia | 0 | 0 | 0 | 100 | 0 |
| Stepping_stone | 0 | 2 | 0 | 0 | 98 |
| ${}^{Ar+He+{}^{1}D}{SS}$ | | | | | |
|  | Hier_Island | Hier_stepping stone | Island | Panmixia | Stepping_stone |
| Hier_Island | 99 | 0 | 0 | 0 | 0 |
| Hier_steppingstone | 0 | 98 | 0 | 0 | 2 |
| Island | 1 | 0 | 100 | 0 | 0 |
| Panmixia | 0 | 0 | 0 | 100 | 0 |
| Stepping_stone | 0 | 2 | 0 | 0 | 98 |
| ${}^{Ar+H+He+{}^{1}D}{SS}$ | | | | | |
|  | Hier_Island | Hier_stepping stone | Island | Panmixia | Stepping_stone |
| Hier_Island | 99 | 0 | 0 | 0 | 0 |
| Hier_steppingstone | 0 | 97 | 0 | 0 | 3 |
| Island | 1 | 0 | 100 | 0 | 0 |
| Panmixia | 0 | 0 | 0 | 100 | 0 |
| Stepping_stone | 0 | 3 | 0 | 0 | 97 |

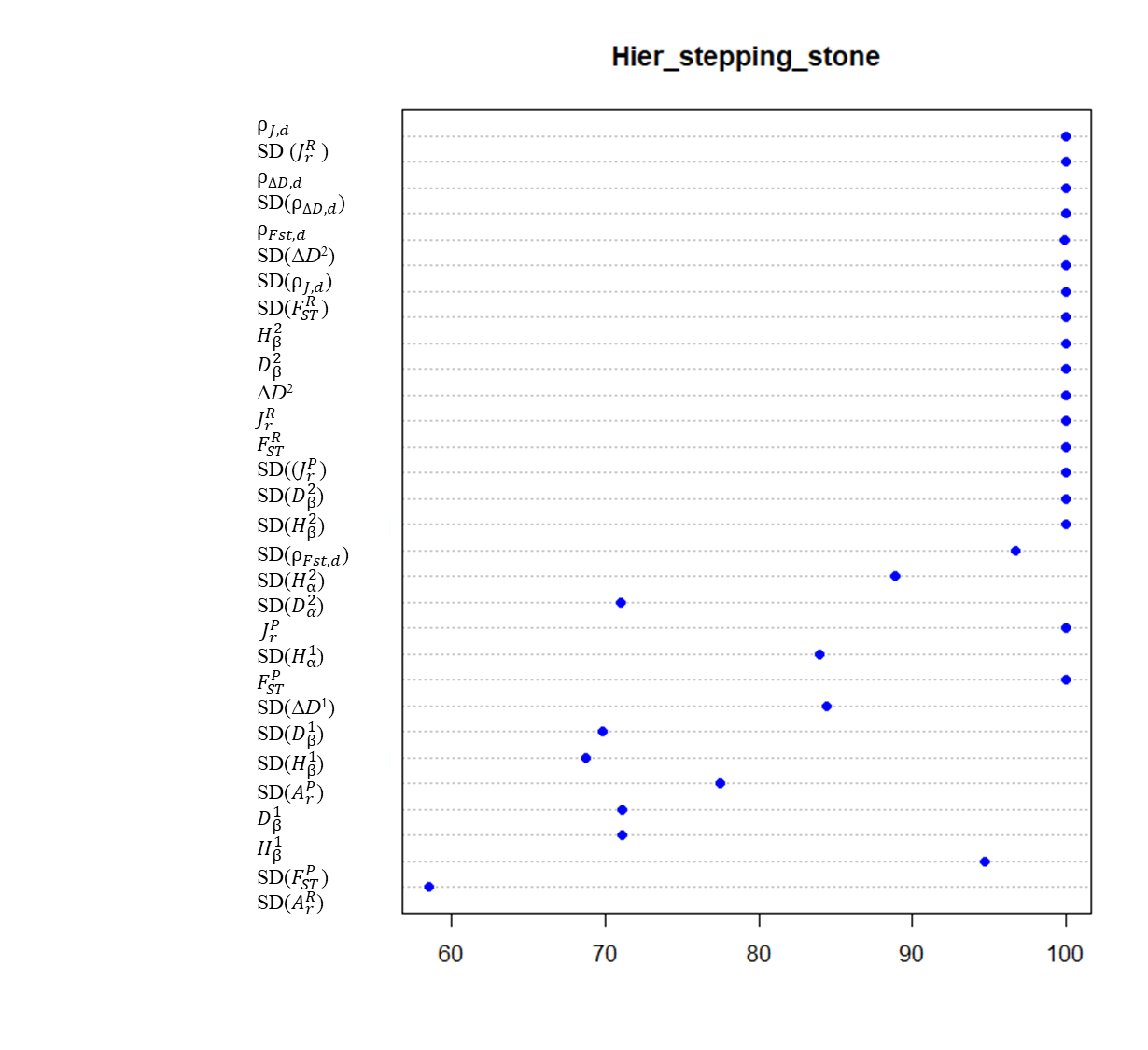

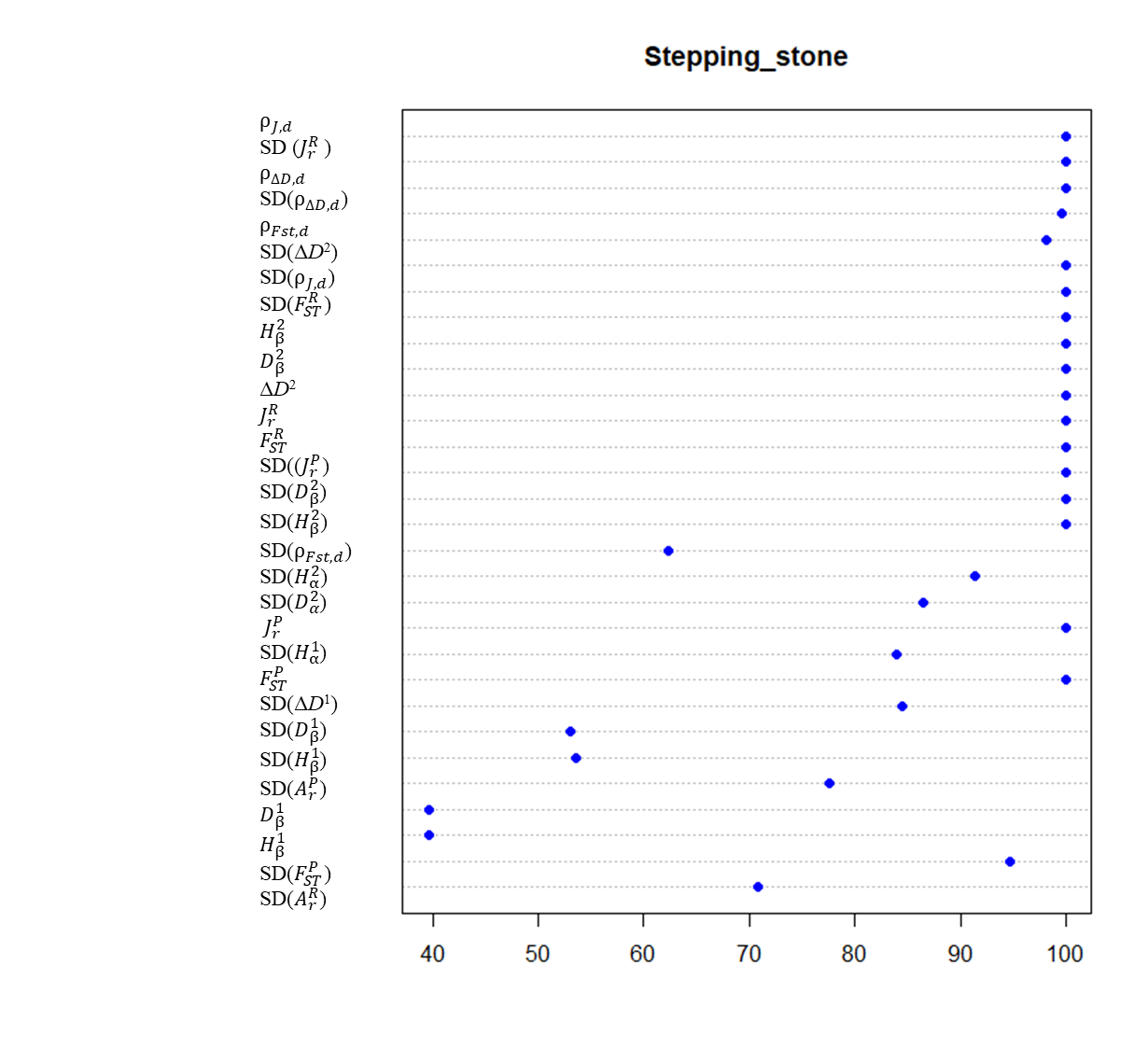

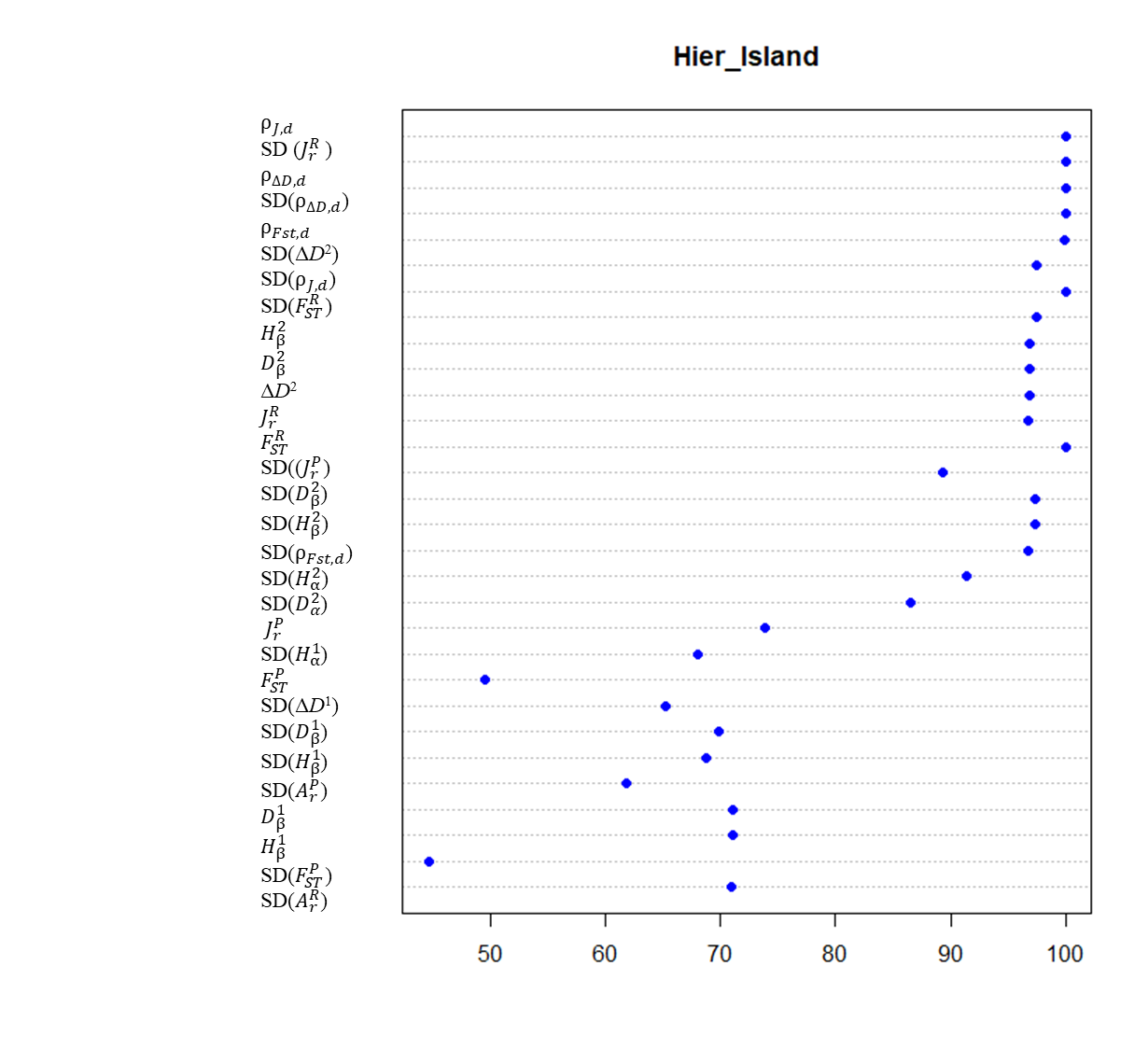

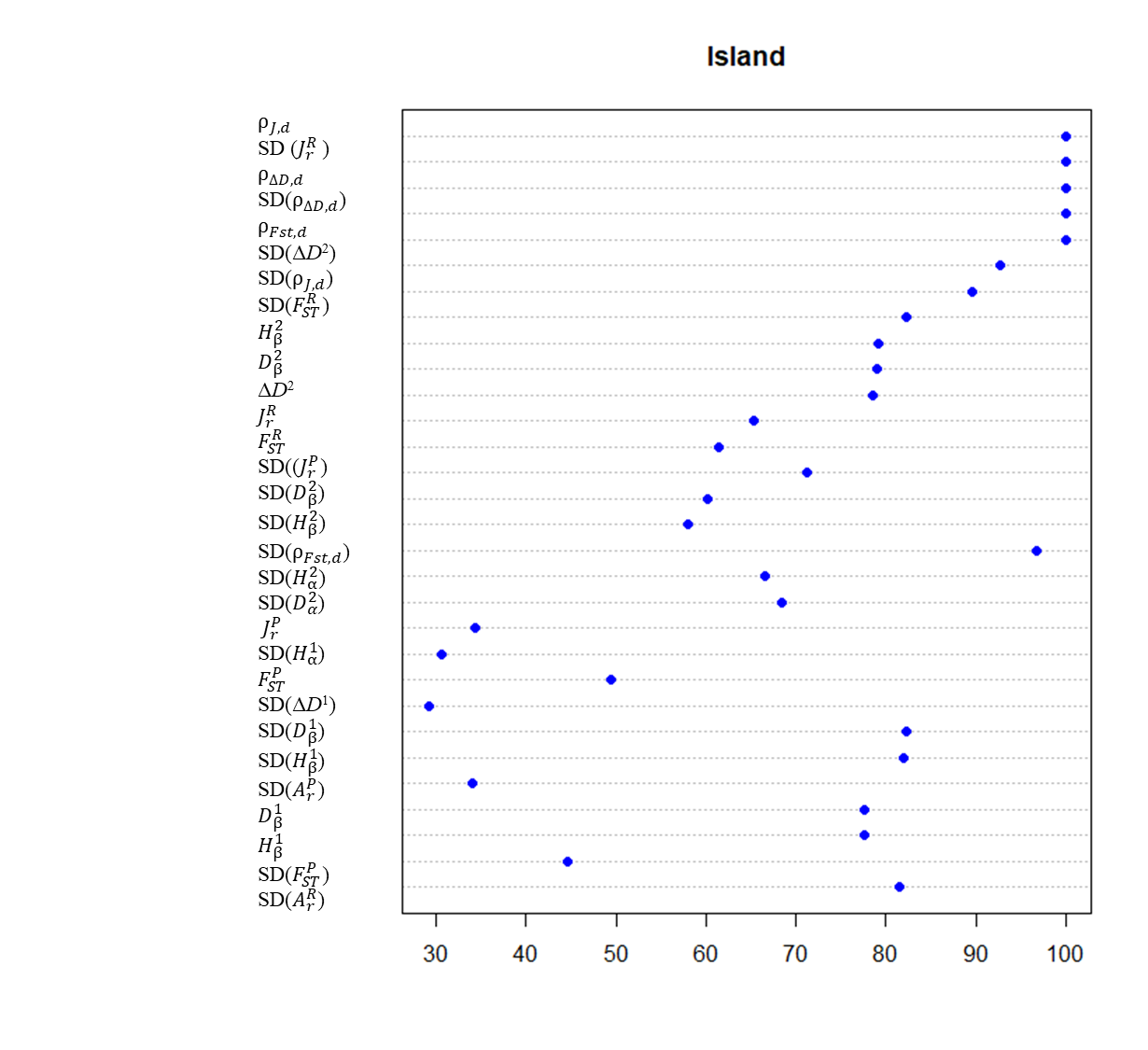

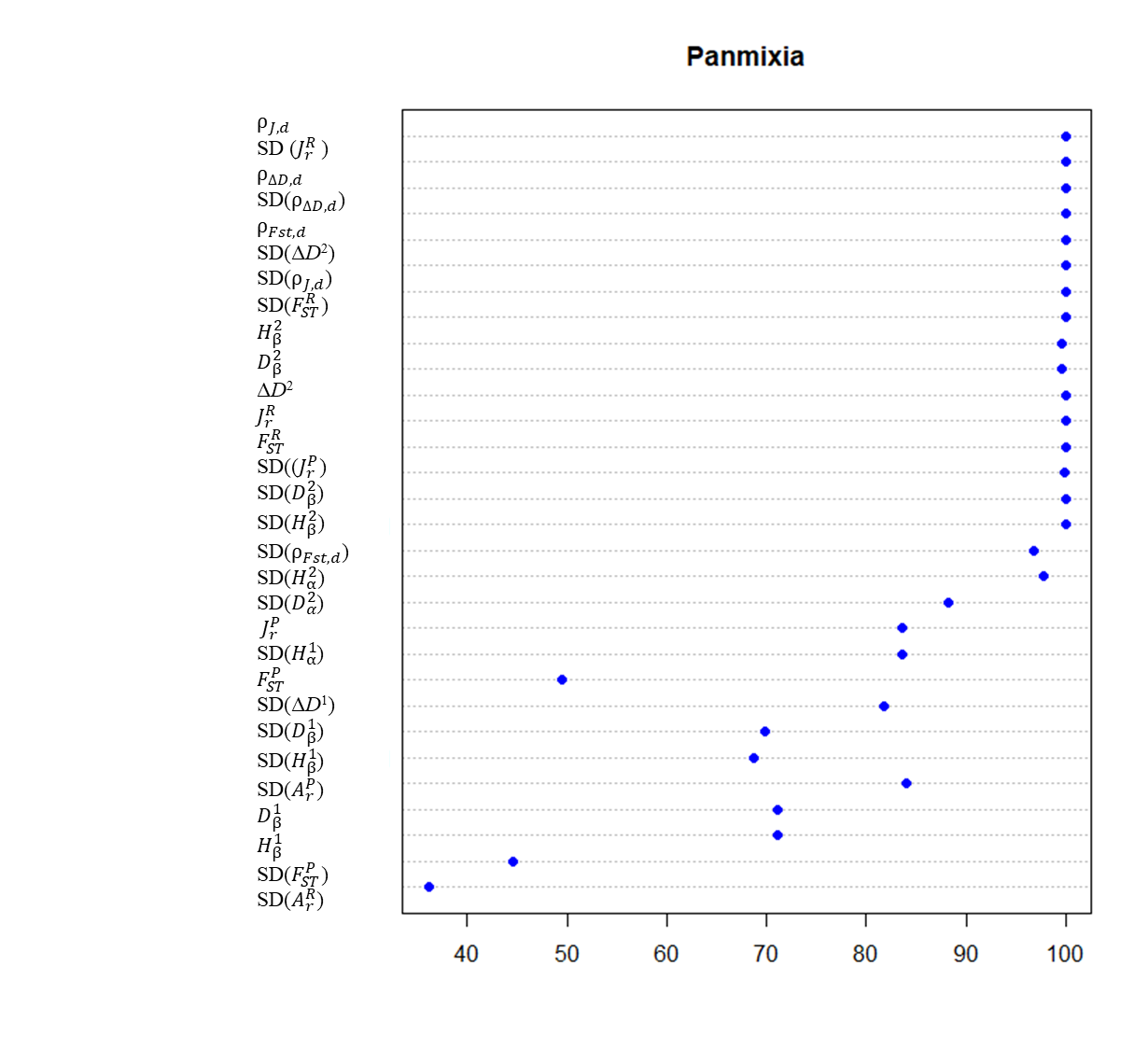

Fig. S1. Scenario-specific variables importance tested from deep neural networks. The ranking is in accordance with the overall importance ranking in Figure 3.

##### Supplementary Methods

##### Model performance metrics

We used confusion matrix, overall performance across all spatial structure scenarios and scenario-specific performance’ of each class of summary statistic to evaluate the power of different sets of summary statistics to discriminate five spatial structure scenarios. The confusion matrix is a cross-tabulation used to describe the performance of a classification model on predicted scenarios given the true scenarios. Table S5 shows an example of confusion matrix which indicates if the model is confusing two scenarios (i.e., commonly mislabeling one as another for two classes, positive vs. negative class). The value A, D represent the number or percentage of a case that is correctly predicted by a model, while B, C represent the number or percentage of a case that is incorrectly predicted by a model.

**Table S5**. Example of confusion matrix of two classes

|  | Target | |
| --- | --- | --- |
| Model | Positive | Negative |
| Positive | A | B |
| Negative | C | D |

Model accuracy (Acc), and unweighted Kappa are the main overall statistics to evaluate the performance of a model. Model accuracy is estimated as the ratio of the total number of correctly predicted scenarios divided by the total number of the training scenarios.

Cohen's kappa coefficient (*κ*) (McHugh 2012) measures inter-rater agreement for classes, defined by,

*κ*= 1- $\frac{{1-p}_{o}}{{1-p}_{e}}$ ,	(S1)

where *p_o_* is the accuracy of correct classification ((A + D)/(A + B+ C+ D)) and is used as the relative observed agreement among scenario here. *p_e_* is the hypothetical probability of random agreement. *p_e_*= ((A + B)/(A + B+ C+ D)) *((A + C)/(A + B+ C+ D)) + ((C + D)/(A + B+ C+ D)) * ((B + D)/(A + B+ C+ D)). If the accuracies are in complete agreement then *κ* = 1. If there is no agreement among the accuracies, *κ* = 0.

For more than two classes, these results are calculated comparing each factor level to the remaining levels (i.e., a "one versus all" approach).
